# A reference model for the combination of an arbitrary number of drugs: A generalization of the Bliss independence model

**DOI:** 10.1101/630616

**Authors:** Wim De Mulder, Martin Kuiper, Åsmund Flobak

## Abstract

It is commonplace to determine the effectiveness of the combination of drugs by comparing the observed effects to a reference model that describes the combined effect under the assumption that the drugs do not interact. Depending on what is to be understood by non-interacting behavior, several reference models have been developed in the literature. One of them is the celebrated Bliss independence model, which assimilates non-interaction with statistical independence. Intuitively, this requires the dose-response curves to have zero as minimal effect and one as maximal effect, a restriction that was indeed adopted by Bliss. However, we show how non-interaction can be interpreted in terms of statistical independence, while nevertheless allowing arbitrary values for the minimal and the maximal effect. Furthermore, our reference model allows the maximal effects of the dose-response curves to be different. In a first step, we construct a basic reference model for the case of two drugs and where the maximal effects of the two individual dose-response curves are assumed to be equal. By relying on the notion of non-interaction in terms of statistical independence, and by introducing two consistency principles, we show how a unique reference model can be derived. In a second step, a more general reference model, allowing the maximal effects to be different while still restricting to two drugs, is then easily constructed from the basic reference model. Finally, an induction step is applied to generalize the reference model to the case of an arbitrary number of drugs, allowing each dose-response curve to have a possibly different maximal effect. Although the minimal effect of the dose-response curves are restricted to be equal, which we show to be a necessary consequence of consistency rules, its value is arbitrary.

**Author summary:** The Bliss independence model is a very popular reference model for drug combinations, meaning that it predicts the combined effect of doses of given drugs under the assumption of non-interaction between these drugs. However, because Bliss described non-interaction as statistical independence, he thought that he had to assume that the minimal effect of all dose-response curves are zero, while the maximal effect of all dose-response curves are one. While it is acceptable that all dose-response curves have minimal effect zero, because this amounts to having a common reference state (i.e. the response when no drug at all is given), it is a severe restriction to force all dose-response curves to have maximal effect one. On the other hand, the Bliss independence model has the advantage that it relies on sound statistical theory, and the assimilation of non-interaction with statistical independence is rather intuitive. We have extended the Bliss independence model to allow the involved dose-response curves to have different maximal effects. This has been done in a rigorous way, where the statistical underlying theory that was used by Bliss remains essentially intact.

## 1 Introduction

### 1.1 Background and goal of the paper

For certain diseases it has become standard practice to administer a combination of drugs instead of a single agent, stimulated by advances in omics and cell biology (Keith, Borisy and Stockwell 2005). Examples of diseases where one has become convinced of the superiority of combining drugs include cancer (Al-Lazikani, Banerji and Workman 2012; Bukowska, Gajek and Marczak 2015; Preuer et al. 2018), Parkinson’s disease (Bitner et al. 2015; Hajj et al 2015; Matsunaga, Kishi and Iwata N 2017; Przuntek et al 1992), HIV (Cihlar and Fordyce 2016; Scourfield, Waters and Nelson 2011; Sombogaard 2018; Wilson, Gallant and Mayer 2009) and asthma (Breton et al 2007; Nelson 2001; Pierarch 2001; Ruggeri et al 2012; Saleh 2008).

However, there is no a priori ground to assume that the combination of individual drugs will result in a larger desired effect than the additive effect of the single drugs. The additive effect in this context is, loosely speaking, to be interpreted as the effect that one expects to result from the combination of the drugs based solely on knowledge of the effects of the single drugs. For a given set of drugs there are basically three possibilities: 1. the drugs do not interact, meaning that no superior nor inferior effect arises by combining them; 2. the drugs do interact and thereby reinforce each other, resulting in a superior effect compared to the additive effect, a phenomenon known as synergy; 3. the drugs are interacting but their combination results in an effect that is inferior to the additive effect of the single drugs, known as antagonism (Bell 2004; Tallarida 2001). Determining which of the three possibilities applies in a particular case relies on having a theoretical model that describes the additive effect of the single agents, referred to as the reference model. Having constructed a model that represents the additive effect, the observed effect can then be compared to this additive effect to determine which of the three aforementioned possibilities hold. It should not come as a surprise that currently there does not exist a universally accepted reference model. The reason is that the construction of a reference model depends on the meaning of the concept of non-interaction between drugs, since non-interaction is the key ingredient in determining the additive effect. Indeed, any interaction between drugs will produce an effect that was not expected based on the effects of the individual drugs. As yet, there does not exist a standard interpretation of non-interaction in the context of drug combinations (Foucquier and Guedj 2015).

In this paper we extend a reference model that is still very popular, despite the fact that it was published a long time ago. The Bliss independence model (Bliss 1939) was published in 1939, and relies on the statistical concept of independence. We also describe non-interaction between drugs as statistical independence between suitably defined random variables. Our model also shows some similarities to the zero interaction potency (ZIP) model (Yadav et al. 2015), a recent model that has received attention from the drug combinations community. More specifically, we rely on the authors’ idea to construct a reference model by fixing a certain drug and describing what happens when a given amount of another drug is added to it.

### 1.2 Outline of the paper

The paper is outlined as follows. Section 2 introduces some notation and describes the assumptions that will be maintained throughout the paper, such as an equal minimal effect for all considered dose-response curves. The Bliss independence model and the ZIP model, which is related work to our research, are briefly reviewed in Section 3. Bliss provided a rather ad hoc similarity between the behavior of drugs and statistical independence, establishing the relationship between probability and dose-response curves by restricting the minimal effect of a dose-response curve to zero and its maximal effect to one. We introduce a sound relationship between a given dose-response curve and the expected value of a properly defined random variable. This establishes an appropriate statistical setting that is equivalent to the deterministic setting that is traditionally considered in describing the effect of a drug for a given dose. Furthermore, the relationship allows the dose-response curve to have arbitrary minimal and maximal effect. The probabilistic setting is outlined in Section 4. In Section 5 we construct our basic reference model, which applies to the combination of two drugs that have the same maximal effect. It is shown how the basic reference model is uniquely identified from the two consistency principles that are introduced in the same section. The basic reference model is then extended, in Section 6, to dose-response curves with a possibly different maximal effect, while still restricting to the case of two drugs. The ideas underlying the construction of a reference model for two drugs are also applicable in constructing a reference model that describes the effect of the combination of an arbitrary number of non-interacting drugs, where each drug has a dose-response curve with a possibly different maximal effect. Induction does the trick, and the general reference model is described in Section 7. Finally, in Section 8 we illustrate our model, thereby indicating the importance of allowing for different maximal effects.

## 2 Notational conventions and assumptions

The considered drugs are conveniently referred to as the first drug, the second drug, etcetera. The dose-response curve of each drug is assumed to be given and is denoted by *f*_*i*_, *i* ∈ ℕ. An arbitrary dose of the *i*th drug is denoted by *x*_*i*_ with corresponding effect given by *y*_*i*_ = *f*_*i*_(*x*_*i*_). We adopt the common assumption that *f*_*i*_ is strictly increasing. At dose 0 each drug obtains its minimal effect *A*_*i*_ = *f*_*i*_(0). However, we assume throughout the paper that the minimal effect of all involved individual dose-response curves are equal to the same value *A* ≡ *A*_*i*_. One can also consider this a logical or commonsense restriction, since not applying a certain drug (which is the same as administering a dose 0 of that drug) should result in the same effect as not applying any other drug. Indeed, in both cases no action has been taken. In terms of a thought experiment, a patient is not able to distinguish between having administered a dose 0 of a certain drug or a dose 0 of any other drug. In other words, logic and commonsense require a unique reference state. However, this argument breaks down in case the drugs are administered at different times, since the influence of other factors might change the reference state over time. But then consistency requires that there exists a data transformation such that the effects of both drugs can be described with respect to a common reference state. This is very similar to the arbitrary choice of a single origin in describing certain physical systems in Euclidean space. The dose-response curves are bounded above by their maximal effect *B*_*i*_, in the sense that 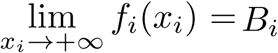. Since, in the case of two drugs, a reference model is meant to give us the additive effect for the combination of a dose *x*_1_ of the first drug with a dose *x*_2_ of the second drug, the reference model is conveniently denoted by *f*_1,2_(*x*_1_, *x*_2_). In the more general case of *n* drugs, the reference model is denoted by *f*_1,…,*n*_(*x*_1_, …, *x*_*n*_).

## 3 Related work

### 3.1 Bliss independence model

For convenience in terms of notation, we review the Bliss independence model for the case of two drugs. The Bliss independence model assumes that *A* = 0 and *B* ≡ *B*_1_ = *B*_2_ = 1. Together with the assumption that the dose-response curves are strictly increasing, it follows that *f*_*i*_(*x*_*i*_) can be interpreted as a probability. This interpretation allows to formulate a reference model in a probabilistic setting, in the following way. Let a patient be given a dose *x*_*i*_ of the *i*th drug, and assume that she either encounters an effect (success) or no effect (failure). Instead of modelling the response in terms of the given dose, the probability of an effect can then be considered. Both are equivalent modelling perspectives, because the given assumptions imply that *f*_*i*_(*x*_*i*_) represents both the effect at dose *x*_*i*_ (in the deterministic setting) and the probability of an effect at dose *x*_*i*_ (in the probabilistic setting). The probabilistic interpretation can be made more explicit by introducing the event *D*_*i*_(*x*_*i*_) that the *i*th drug results in an effect at dose *x*_*i*_. By the given arguments it then holds that the probability of this event is given by *f*_*i*_(*x*_*i*_) = *P* (*D*_*i*_(*x*_*i*_)). Bliss then defines non-interaction between the two drugs as the statistical independence of the associated events *D*_1_(*x*_1_) and *D*_2_(*x*_2_), implying that

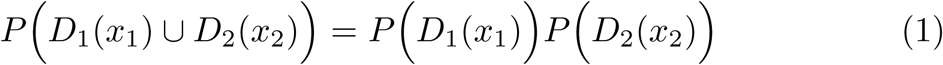

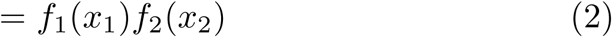

Continuing the reasoning in terms of probability, the effect of the combination of dose *x*_1_ of the first drug with dose *x*_2_ of the second drug, is then interpreted as the probability that at least one of the events *D*_1_(*x*_1_) and *D*_2_(*x*_2_) occurs. Indeed, an effect will be observed when combining the two drugs if at least one of the drugs results in an effect. Using the non-interaction principle given by (1), and the probabilistic formulation of the probability of the union of two events, the Bliss independence model is then given by

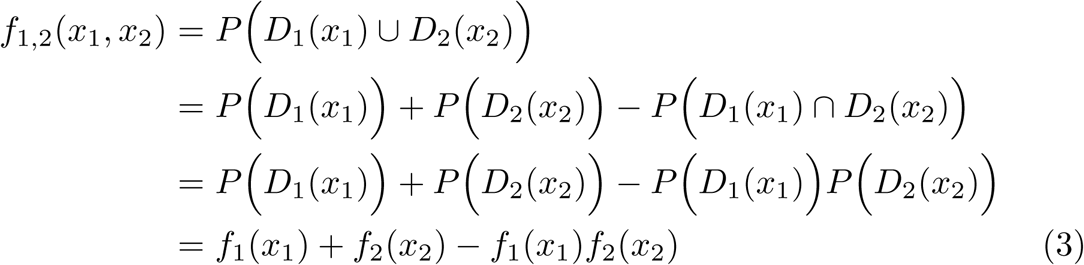

### 3.2 Zero interaction potency

The zero interaction potency (ZIP) model applies to the case of two drugs. Furthermore, the authors assume a Hill curve as dose-response curve, and that the maximal effects are equal: *B* ≡ *B*_1_ = *B*_2_. This results in the following description of the individual dose-response curves:

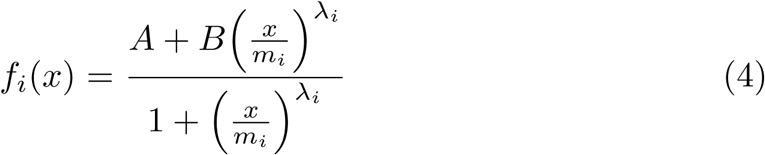

with *m*_*i*_ and *λ*_*i*_ parameters.

As a first step, the authors assume that *A* = 0 and *B* = 1. The idea is then to consider the addition of a dose *x*_2_ to a dose *x*_1_ of the first drug. Thus the first drug is given the focus, and the question is how the effect of the first drug changes when a certain dose of the second drug is added to it. Non-interaction according to the authors means that the effect of this combination can still be represented by a curve similar to the Hill curve (4) with *i* = 1 (i.e. the dose-response curve for the first drug). The parameters *m*_1_ and *λ*_1_ are not influenced by this combination, and thus retain their original values as before the combination. Otherwise, the shape of the curve would be changed, implying that the second drug would have changed the effectiveness of the first drug. This would imply interacting behavior, contrary to the assumption that the drugs do not interact. However, the minimal effect of the dose-response curve of the first drug will be changed from *A* = 0 to *A* = *f*_2_(*x*_2_), since adding a dose *x*_2_ of the second drug to a dose 0 of the first drug results, of course, in the effect *f*_2_(*x*_2_). Denoting the effect that results by adding a dose *x*_2_ of the second drug to a dose *x*_1_ of the first drug by 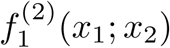 we then have that

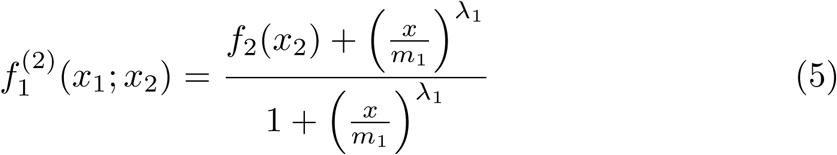

They go on by showing that the conceptual view of adding the second drug to the first drug is equivalent to adding the first drug to the second one, i.e. 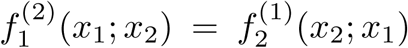. The model is thus consistent with drug labelling, meaning that it does not matter which drug is referred to as the first drug. Since both formulations are the same, this can be taken as the reference model. By applying some algebra, they show that 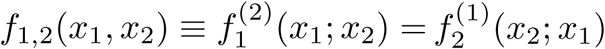 can be written as:

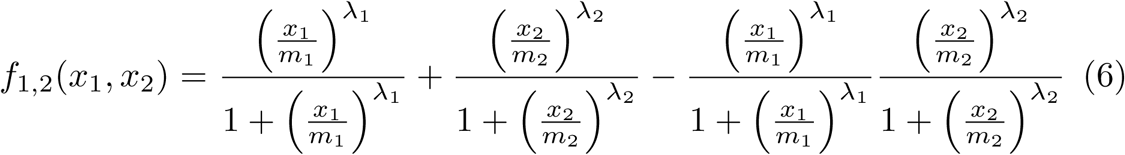

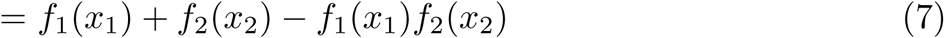

which is the same as the Bliss independence model, except for the fact that the authors restrict the dose-response curves to Hill curves. In the special case where *A* = 0 and *B* = 1 the ZIP model is thus more restrictive than the Bliss independence model.

In a second step they allow that *A* and *B* are arbitrary values. It is, however, without motivation that they state their reference model for this more general case:

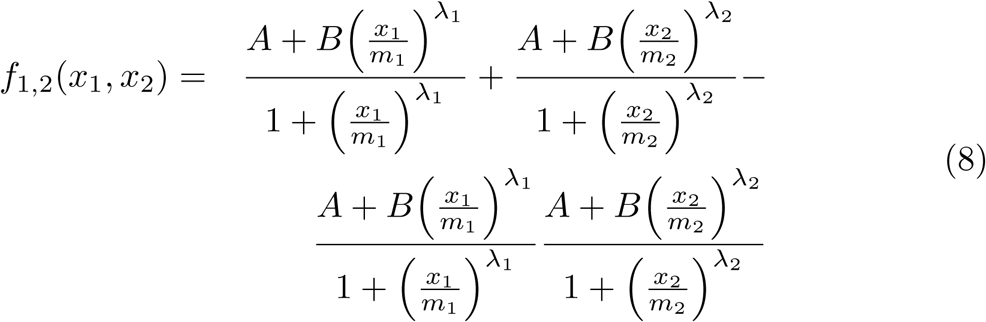

Seemingly, the authors combined the expressions in (4) and (6) to make an educated guess on the proper position of *A* and *B* when the more general case is considered.

## 4 Introduction of a general probabilistic setting

We make use of the idea of thinking in terms of a probabilistic setting, as in the Bliss independence model, although we do not restrict the minimal and maximal effects. This is done by separating, in a certain sense, the minimal and maximal effect from the probabilistic part in the description of a dose-response curve.

### 4.1 Alternative description of a dose-response curve

We introduce the following very simple, yet extremely useful theorem.

#### Theorem 1

*Any function f* (*x*), *with x* ≥ 0, *that is continuous and strictly increasing, with f* (0) = *A and* 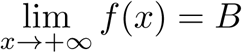, *can be written as*

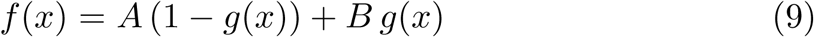

*with g*(*x*) *a function that is continuous and strictly increasing, and for which g*(0) = 0 *and* 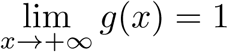.

*Proof Define* 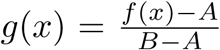 *if A < B, which clearly has the required properties. Then it follows that f* (*x*) = *A* (1 −*g*(*x*))+ *B g*(*x*). *When A* = *B, it follows that f* (*x*) = *A, in which case f can still be written as (9) and this for any g that has the required properties.*

Since the function *f* in the theorem has exactly the properties that we require for an individual dose-response curve (cf. Section 2), we can thus as well define a dose-response curve in the following way.

#### Definition 1

A dose-response curve *f* (*x*) is a function of the form

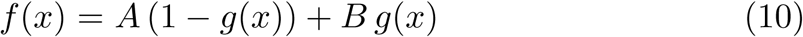

with *g*(*x*) a function that has the following properties:

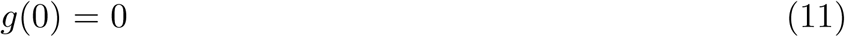

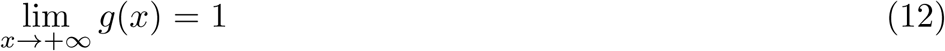

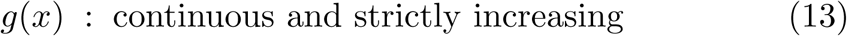

Observe that (10) expresses a dose-response curve as a weighted combination of the minimal effect and the maximal effect, the weights being determined by the function *g*.

We introduce a convenient shorthand notation for the expression in (10).

#### Definition 2

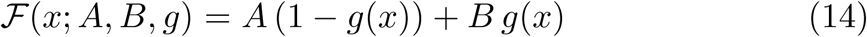

### 4.2 Probabilistic interpretation of the dose-response curve

The description *f* (*x*) = *A*(1 − *g*(*x*)) + *Bg*(*x*) for the response of a drug can be given an elegant interpretation. Notice that (11)-(13) implies that 0 ≤ *g*(*x*) ≤ 1. This means that *g*(*x*) for any fixed dose *x* can be interpreted as a probability. This is similar to the Bliss independence model, with the important difference that in our formulation it is not the dose-response curve itself that has the meaning of a probability (which would require the minimal effect to be zero, and the maximal effect to be one), but only a certain part of it, namely *g*. Again similar to the rationale behind the Bliss independence model, the probabilistic setting implies that a patient who is given a dose *x* of a drug either encounters an effect (success) or no effect (failure). This happens at random, with the probability of success given by *g*(*x*) and the probability of failure by 1 − *g*(*x*). Furthermore, when there is an effect, we consider the effect to be equal to the maximal effect *B*, and the absence of an effect is represented by *A* (since the minimal effect results from a dose 0, i.e. *A* corresponds to a rest state). We denote the random variable in this probabilistic setting by *Z*(*x*):

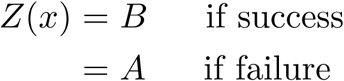

with

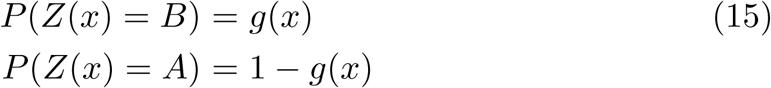

According to the definition of expectation, the expected value of *Z*(*x*) is given by

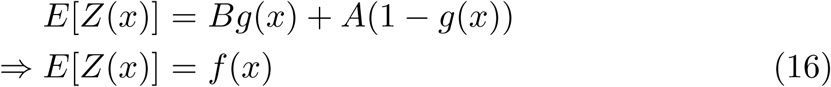

using (10). The drug that is used in the deterministic case, where a dose *x* generates a definite effect *f* (*x*) in any given patient, could be referred to as the deterministic drug. The drug considered in the probabilistic setting, where a dose *x* either generates effect *A* or effect *B* could then be referred to as the probabilistic drug. Equation (16) then expresses a duality between the deterministic drug and the probabilistic drug: the expected value of the effect of the probabilistic drug equals the effect of the deterministic drug. In this sense, the deterministic drug and the probabilistic drug are equivalent.

## 5 Basic reference model: two drugs and *B*_1_ = *B*_2_

As a first step, we assume that there are only two drugs to be combined and that *B* ≡ *B*_1_ = *B*_2_. The reference model that will be constructed for this case is called the basic reference model.

### 5.1 Introduction

Using the general description of a dose-response curve, given by (10), and using the assumption that *B*_1_ = *B*_2_ = *B*, the dose-response curves of the two drugs are described by

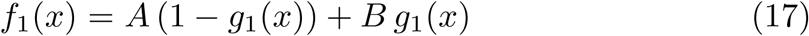

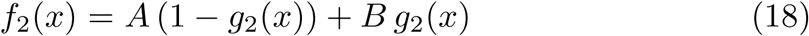

We now rely on one of the ideas that underlies the ZIP model: constructing a reference model by taking the point of view that one of the drugs is added to the other one, assuming that the reference model will retain certain properties of the dose-response curve of the drug to which the other one is added. Let us first consider the case where a dose *x*_2_ of the second drug is added to a dose *x*_1_ of a first drug. As in Section 3.2, the resulting effect is denoted as 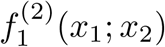. Since this function describes the response for given doses, similar to the doseresponse curve for a single drug, the expression for 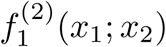 should conform the description of a general dose-response curve given by (10). However, the notation in (10) is to be generalized to take two doses into account, each dose corresponding to a different drug. This can be done as follows:

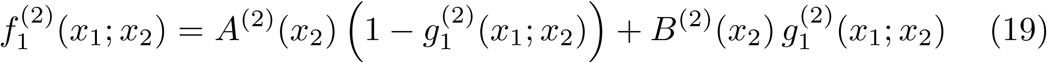

where *A*^(2)^(*x*_2_) and *B*^(2)^(*x*_2_) are the new minimal and maximal effects after a dose *x*_2_ of the second drug is added to any dose *x*_1_ of the first drug. The reason that we consider the new minimal and maximal effects to be functions of *x*_2_ is simply that it is possible that these quantities depend on the exact dose *x*_2_ that is added. These quantities are as yet unknown. Furthermore, reasoning in terms of the probabilistic setting outlined in Section 4, results in the generalization of *g*_1_(*x*_1_) to 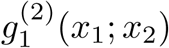. Whereas *g*_1_(*x*_1_) is the probability that dose *x*_1_ of the first drug results in an effect, 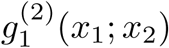 expresses the probability that an effect is observed after a dose *x*_2_ of the second drug is added to a dose *x*_1_ of the first drug. Following Bliss, we define non-interaction as statistical independence between the random variables *Z*_1_(*x*_1_) and *Z*_2_(*x*_2_), implying that

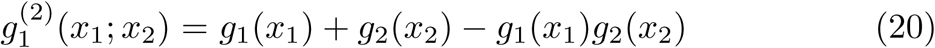

Consider now the other point of view: adding a dose *x*_1_ of the first drug to a dose *x*_2_ of the second drug. The general description of the combined effect is simply obtained from (19) by changing indices. This also applies to 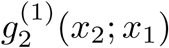, which is obtained from (20) by changing indices. It is easily seen that this results in the same expression as for 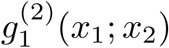, given by the right hand side of (20). The right hand side of (20) can therefore be taken as the probability of observing an effect when a dose *x*_1_ of the first drug is combined with a dose *x*_2_ of the second drug. That is, we do not need to consider which drug is added to the other one in describing the probability of a combined effect. We may thus introduce the notation *g*_1,2_(*x*_1_, *x*_2_) to represent this probability:

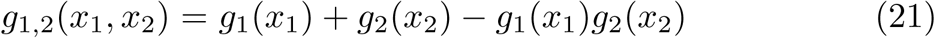

It remains to determine the unknowns *A*^(2)^(*x*_2_) and *B*^(2)^(*x*_2_). We determine these functions by introducing two consistency principles, i.e. principles that should be obeyed by any reference model because of logical consistency.

### 5.2 First consistency principle and application

The combined effect by adding a dose 0 of the second drug to a dose *x*_1_ of the first drug is, logically speaking, the same as the effect of dose *x*_1_ of the first drug. In other words, administering a dose 0 of the second drug is the same as not administering that drug. The same reasoning applies, of course, when a dose 0 of the first drug is added to a dose *x*_2_ of the second drug. This explains the following logical principle.

#### Definition 3

First consistency principle

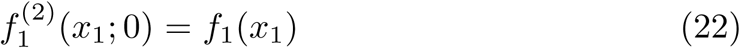

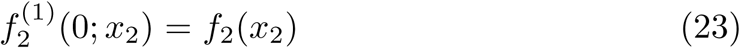

The first consistency principle allows to determine *A*^(1)^(0), *A*^(2)^(0), *B*^(1)^(0) and *B*^(2)^(0).

#### Theorem 2

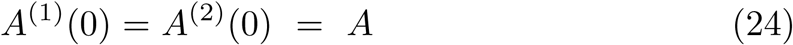

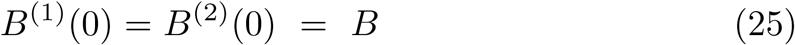

*Proof* If we apply (22) to (19) and (17) we find:

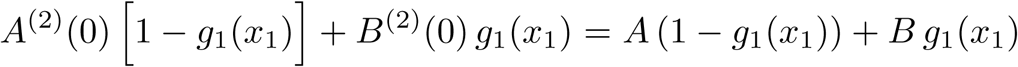

where we used the expression for 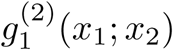 given by (20). If we choose *x*_1_ = 0 in the above equality we find that

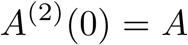

In the same equality, taking on both sides the limit *x*_1_ *→* + *∞* and using (12) results in

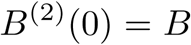

By the same reasoning we find that *A*^(1)^(0) and *B*^(1)^(0) are as promised by the theorem.

### 5.3 Second consistency principle and application

The thought experiment of adding the second drug to the first drug should result in the same combined effect as adding the first drug to the second one. That is, the labels ‘first drug’ and ‘second drug’ should be arbitrary.

#### Definition 4

Second consistency principle

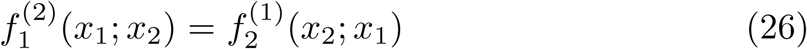

Notice that the first and the second principle together imply the equality of the minimal effects of both dose-response curves, an assumption which was made in Section 2. It now turns out that this is not merely a convenient or commonsense assumption, but rather a restriction dictated by consistency. Indeed, applying the two consistency principles given by (22)-(23) and (26) to the case *x*_1_ = *x*_2_ = 0, we find:

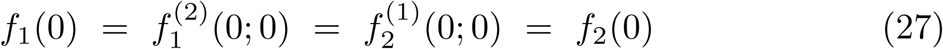

The second principle can be used to completely identify the unknown functions *A*^(1)^(*x*_1_), *B*^(1)^(*x*_1_), *A*^(2)^(*x*_2_) and *B*^(2)^(*x*_2_).

#### Theorem 3

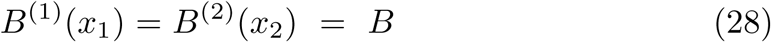

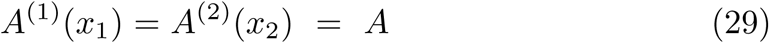

*Proof By (19) and (21) the second consistency principle is equivalent to*

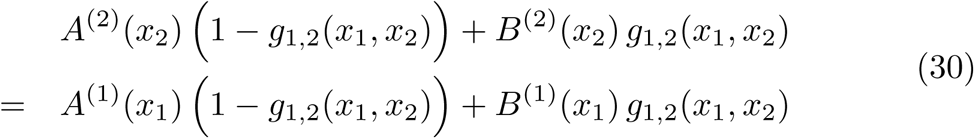

*Taking on both sides the limit x*_1_ *→* +*∞ results in:*

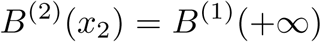

*where B*^(1)^(+*∞*) *is shorthand for 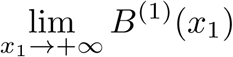. The above equality holds for all x*_2_, *in particular for x*_2_ = 0. *But Theorem 2 tells us that B*^(2)^(0) = *B, cf. (25). Thus*

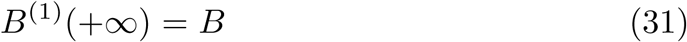

*Since a non-interacting drug cannot result in antagonistic effects, adding a lower dose of the first drug to the second drug cannot result in a higher maximal effect of the combination than adding a higher dose of the first drug to the second drug. That is to say, 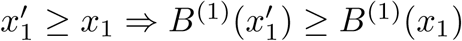, and therefore*

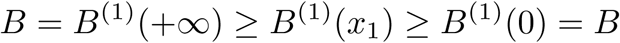

*where we used (25) and (31). Consequently:*

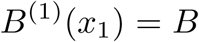

*and by the same reasoning:*

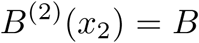

*Thus we have found that B*^(1)^(*x*_1_) = *B*^(2)^(*x*_2_) = *B. If we plug this into equality (30) we see that A*^(2)^(*x*_2_) = *A*^(1)^(*x*_1_). *This holds for all x*_1_, *x*_2_, *in particular for x*_2_ = 0. *By (24) this implies that A*^(1)^(*x*_1_) = *A. The same reasoning results in A*^(2)^(*x*_2_) = *A.*

### 5.4 Reference model

We can now use the values for *B*^(1)^(*x*_1_) and *A*^(1)^(*x*_1_), given by (28) and (29), in the description of 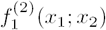, as given by (19). We thus have found our reference model as

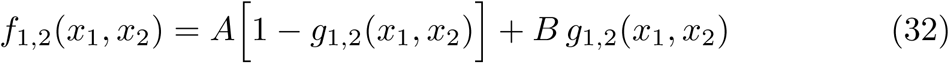

The second consistency principle required that 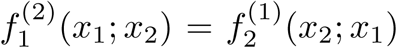. It is easily seen that the above expression for the reference model is indeed independent of the choice of labelling, i.e. *f*_1,2_(*x*_1_, *x*_2_) = *f*_2,1_(*x*_2_, *x*_1_).

As an intermediate conclusion, it is useful to summarize how this reference model was obtained. We started from the common assumption that the individual dose-response curves are increasing, obtaining a minimal effect at dose 0 and a maximal effect at an infinite dose. Then we managed, using the apparently obvious theorem 1, to separate the minimal and maximal effect from the remaining part of the dose-response curve. For that remaining part, i.e. *g*_*i*_(*x*_*i*_), *i* = 1, 2, it was shown how it can be interpreted as the probability that the maximal effect is observed for a given dose *x*_*i*_ for the *i*th drug. The expected value of the associated random variable equals the dose-response curve *f*_*i*_ for dose *x*_*i*_. The dose-response function for the combination of the drugs, i.e. the reference model, was then also assumed to have this structure, although generalized for a combination of doses rather than for a single dose. Two consistency principles and one non-interaction principle (essentially the same one as introduced by Bliss, although applied at the level of *g* instead of at the level of the dose-response function) resulted in the determination of the unknowns in the reference model. It is striking that these three principles are necessary *and* sufficient to completely determine the reference model. That is, one the one hand no further assumptions are needed to completely describe the reference model, and on the other hand the three principles are all needed to uniquely identify the reference model.

### 5.5 Comparison to Bliss independence and ZIP

#### 5.5.1 Comparison to Bliss independence

Our model generalizes the Bliss independence model. To see this, let *A* = 0 and *B* = 1. Then our reference model, given by (32), reduces to

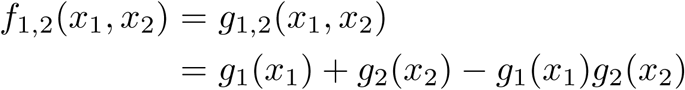

Furthermore, with *A* = 0 and *B* = 1 it follows from (10) that *g*_*i*_ = *f*_*i*_, and thus the above expression becomes:

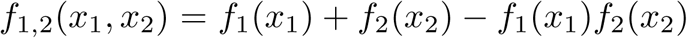

which is equal to the Bliss independence model given by (3).

#### 5.5.2 Comparison to ZIP

Our model generalizes ZIP in the basic case where *A* = 0 and *B*_1_ = *B*_2_ = *B* = 1. This is obvious, as our model generalizes the Bliss independence model, and since the Bliss independence model is itself a generalization of ZIP in the basic case (cf. Section 3.2).

For the more general case where *A* and *B* = *B*_1_ = *B*_2_ are arbitrary, our reference model produces another effect than ZIP. However, we notice that there is a severe inconsistency in the ZIP model in this case. It is clear that it should hold that *f*_1,2_(0, 0) = *A*, since not administering the first drug nor the second drug should have the reference state as resulting effect. However, this is not the effect that is produced by the ZIP model, as seen from (8):

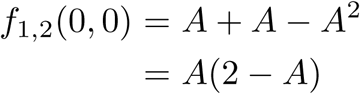

which is different from *A*, unless *A* = 0 or *A* = 1. The inconsistency becomes even more severe if the minimal effect *A* is larger than 2, since in this case the ZIP reference model says that the reference state corresponds to a negative value: *A*(2 - *A*) *<* 0. The same inconsistency applies to the maximal effect. Of course, for effects between the minimal and maximal effect the consistency of the model is not easily verified, as in this case there are no obvious values to compare the value of the reference model to. However, since the reference model is continuous, it is clear that at least effects that are close to the minimal or maximal effect will also be incorrect. It is striking that no single reviewer has taken the time to verify these easy to check consistency conditions.

## 6 More general reference model: two drugs and possibly *B*_1_ *≠ B*_2_

### 6.1 Description

In the basic case, where the two drugs have the same minimal and the same maximal effect, we have found that the reference model is given by

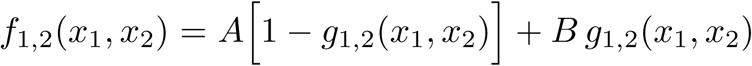

Let us now consider the case where, possibly, *B*_1_ *≠ B*_2_, while still restricting to the case of two drugs. We define, for later use, *β* = min*{B*_1_, *B*_2_*}*. To be definite, and without loss of generality, let *β* = *B*_1_.

Notice that the term *B g*_1,2_(*x*_1_, *x*_2_) in the above expression for *f*_1,2_(*x*_1_, *x*_2_) which applies to the basic case where *B*_1_ = *B*_2_ = *B*, is the term that involves the maximal effect *B*. This term can be written as

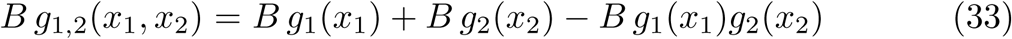

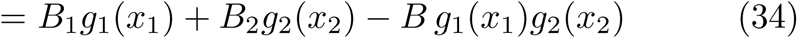

where we used (21). This suggests that in the general case, where *B*_1_ *≠ B*_2_, the term *g*_*i*_(*x*_*i*_) is preceded by the corresponding maximal effect *B*_*i*_. However, it is not clear by which factor the term *g*_1_(*x*_1_)*g*_2_(*x*_2_) should be preceded. We hypothesize that it is a convex combination of *B*_1_ and *B*_2_. This implies that *f*_1,2_(*x*_1_, *x*_2_) in the general case can be written as

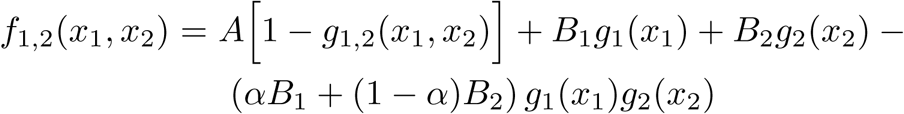

with *α* ∈ [0, 1] unknown. Since *B*_2_ *≥ B*_1_, we should have that a infinite dose of the second drug results in the effect *B*_2_, regardless of the dose *x*_1_ of the first drug:

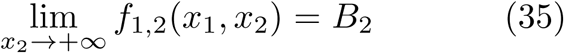

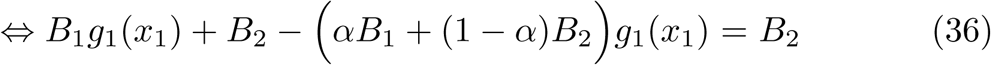

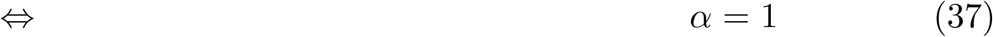

where we made use of 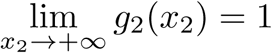. We thus have the following reference model for the case of arbitrary *B*_1_ and *B*_2_:

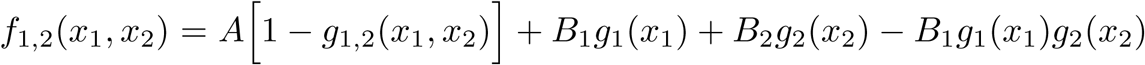

Simple algebra shows that this reference model can also be written as, still assuming that *β* = *B*_1_:

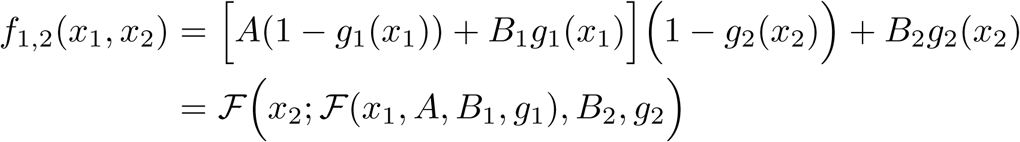

where we used Definition 2 in going to the last line. The reference model thus has the same structure as an individual dose-response curve, as given by (14), although the minimal effect is not a constant anymore, but another doseresponse curve, namely *ℱ* (*x*_1_, *A, B*_1_, *g*_1_) = *f*_1_(*x*_1_). If *β* = *B*_2_, the reference model is simply obtained by suitably changing indices.

This gives the following description of the reference model:

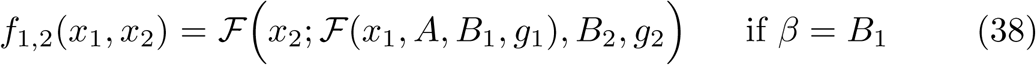

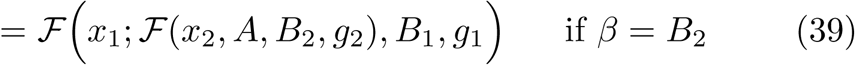

### 6.2 Consistency of the reference model

We derive some properties possessed by the reference model, thereby demonstrating the plausibility and logical consistency of it. We restrict to stating the properties, without performing detailed algebraic derivations, since all properties are obtained by rather straightforward algebra. Furthermore, to be definite, we assume that *β* = *B*_1_ and thus that the reference model is described by (38). The same, or similar, properties can be derived for the case where *β* = *B*_2_.

We have the following:

– The reference model reduces to the basic reference model given by (32) when *B*_1_ = *B*_2_ = *B*.
– The value of the reference model in (0, 0) equals *A*, i.e. *f*_1,2_(0, 0) = *A*. Compare this to the inconsistency that is present in the ZIP model, cf. Section 5.5.2.
– As already established, the maximal effect equals *B*_2_ and is obtained for an infinite dose of the second drug:

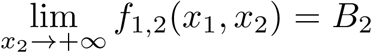
– The model obeys the first consistency principle:

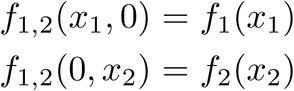

What is missing from the list of properties is the equality *f*_1,2_(*x*_1_, *x*_2_) = *f*_2,1_(*x*_2_, *x*_1_). This is explained by the fact that the maximal effects of the drugs are different, which results in an asymmetry between the drugs. This can be understood as follows. Suppose that a dose *x*_1_ of the first drug is kept fixed, while increasingly larger doses *x*_2_ of the second drug are added. The larger *x*_2_, the closer the effect will be to *B*_2_. Consider now the other direction, where a dose *x*_2_ of the second drug is kept fixed, while increasingly larger doses *x*_1_ of the first drug are added to it. In this case, assuming that *B*_1_ *< B*_2_, the effect can never become arbitrarily close to *B*_2_. Indeed, the first drug has a smaller maximal effect than the second drug and is therefore less effective. While in the case where *B*_1_ = *B*_2_ an infinite dose of the first drug results in the same combined effect as an infinite dose of the second drug, irrespective of the finite dose of the other drug, this principle does not hold in the case where *B*_1_ ≠ *B*_2_ The general reference model shows that

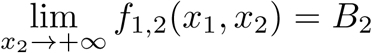

While

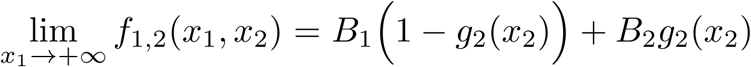

The last equation thus shows that an infinite dose *x*_1_ of the first drug results in an effect that is between *B*_1_ and *B*_2_, depending on the dose *x*_2_ of the second drug. On the other hand, an infinite dose *x*_2_ of the second drug results in the maximal effect *B*_2_, irrespective of the dose *x*_1_ of the first drug.

## 7. General reference model: arbitrary number of drugs and possibly different maximal effects

### 7.1 Considering three drugs

We start our generalization to an arbitrary number of drugs by considering three drugs. We label the drugs such that *B*_1_ *≤ B*_2_ *≤ B*_3_. Let us first combine the first drug and the second drug. The combined effect is, as we know, given by the reference model (38):

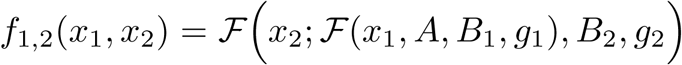

The combination can be considered a new single drug, to which we refer as the zeroth drug for convenience, since the above dose-response function has the same structure as a single-dose response curve, cf. Definition 2. To combine it with the third drug, we notice that the zeroth drug has as maximal effect *B*_2_ (cf. Section 6.2), which is smaller (or possibly equal) to the maximal effect *B*_3_, by assumption. This implies that combining the zeroth drug with the third drug can be performed in the same way as described in Section 6.1 for the case of two single drugs. The resulting reference model *f*_1,2,3_(*x*_1_, *x*_2_, *x*_3_) is then given by

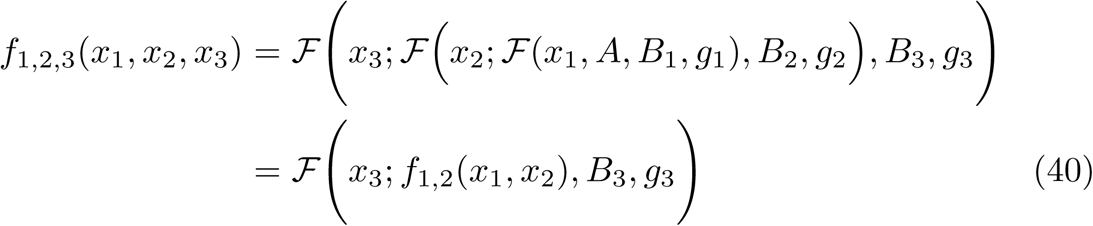

using (38) in going to the last line.

Let us verify the special case *A* = 0, *B*_1_ = *B*_2_ = *B*_3_ = 1. According to Definition 2 we then have that *ℱ* (*x*_1_, *A, B*_1_, *g*_1_) = *g*_1_(*x*_1_). This implies that

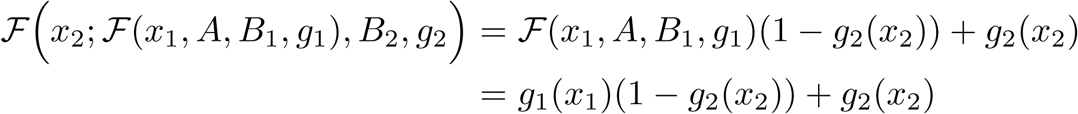

and thus *f*_1,2,3_(*x*_1_, *x*_2_, *x*_3_), given by (40), becomes

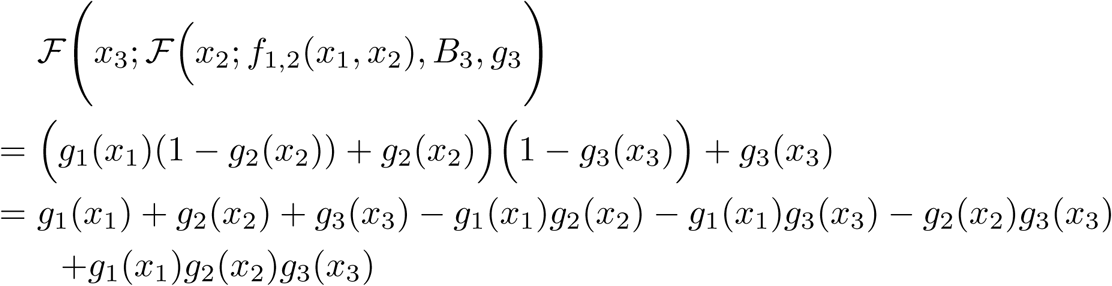

Notice that using (15) the right hand side of the last line equals

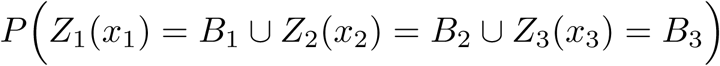

Consequently, our reference model reduces to the Bliss independence model for three drugs in this special case, since the special case implies that *g*_*i*_ = *f*_*i*_ by (10).

### 7.2 Considering *n* drugs

The extension to an arbitrary number of drugs is now straightforward. Let us first introduce some convenient shorthand notation.

#### Definition 5

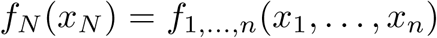

Now label the drugs such that *B*_1_ *≤ B*_2_ *≤ … ≤ B*_*n*_. The reference model for three drugs, given by (40), then suggests the following recursion relation:

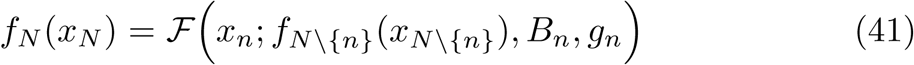

Of course, induction requires to proof that *if f*_*N\{n}*_(*x*_*N\{n}*_) is the reference model for *n -* 1 drugs, *then* the reference model for *n* drugs is indeed given by the above expression. However, this is easily verified by relying on the description of constructing the reference model for two and three drugs above.

Notice the elegance of the formulation of the general reference model, given by (41). Although an arbitrary number of drugs is involved, the description of the reference model is still similar to the dose-response curve of a single drug, cf. (14), although, of course, the parameters are different.

An important strength of the reference model is that it does not assume a specific function for the individual dose-response curves. The involved doseresponse curves are only required to obey the general form of a dose-response curve as established by Definition 1. This definition only contains some mild assumptions on *g*, especially that it is increasing. In particular, it is allowed that the dose-response curves take different functional representations, for example one dose-response curve is a Hill curve while another one assumes an exponential form.

## 8. Illustration

In this section we illustrate the difference between our reference model, to which we will also refer as the unrestricted Bliss reference model (because it extends the original formulation of Bliss’ reference model to allow for arbitrary maximum effects), and the reference model originally developed by Bliss, also referred to as the restricted Bliss reference model (since it assumes that all maximum effects are one). We first generate two artificial data sets that represent dose-responses of two different drugs. For both data sets we consider 50 randomly generated doses *d*_1_, *…*, *d*_50_ in [0,10]. The response of the corresponding drug is assumed to have the following exponential form:

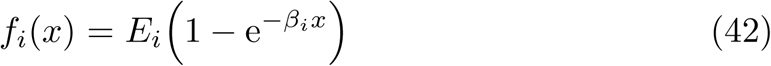

with *i* ∈ *{*1, 2}, and where *β*_*i*_ *>* 0 is a shape parameter. Due to the above representation of the response, it is implicitly assumed that both minimum effects are zero. The maximum effect is given by *E*_*i*_. However, due to measurement errors, we consider the measured responses to be given by

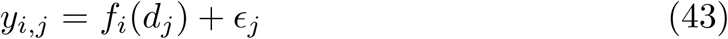

where *y*_*i,j*_, *j* = 1, *…*, 50, refers to the response to dose *d*_*j*_ of the *ith* drug, and where *E*_*j*_ is a random number generated from the normal distribution with mean 0 and variance 0.02. We choose:

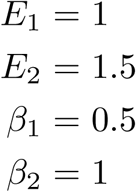

It is important to note that the maximum effects of the two drugs are different. After having generated the two data sets *D*_*i*_ ={(*d*_*i*_, *y*_*i,j*_) | *j* ∈ {1, *…*, 50}}, *i* ∈{1, 2}, we construct a dose-response curve for each data set. We use the exponential form as model. Since the unrestricted reference model allows arbitrary maximum effects, the dose-response curves that will be used for this reference model are given by 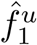 and 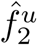:

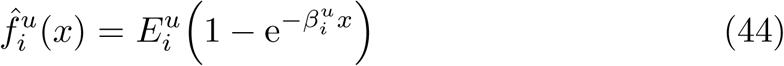

where the parameters 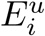 and 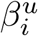 have to be estimated from the corresponding data set *D*_*i*_. In this notation the superscript ‘u’ refers to ‘unrestricted’. We use the function nlsLM from R to estimate these parameters. On the other hand, the dose-response curves that will be used for the restricted reference model need to have maximum effect one. This requires to set *E* = 1 in (42), implying that the dose-response curves to be used for the restricted reference model are modeled by

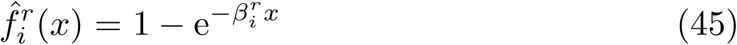

where the single parameter 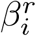 is also estimated using nlsLM in R. Table 1 shows the estimated parameters. Figs. 1 and 2 show the estimated doseresponse curves for the first and second drug, both for the unrestricted model and the restricted model, together with the responses *y*_*i,j*_. Notice that for the first drug the estimated dose-response curves 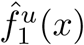 and 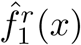 are nearly identical, which is explained by the fact that the maximum effect is one, a case that can be handled in the context of Bliss’ original formulation of drug combinations. However, for the second drug there is a clear difference between 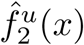 and 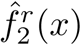, since the dose-response curve to be used for the restricted model, i.e.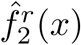, assumes that the maximum effect is one, which does not hold, since *E*_2_ = 1.5

**Table 1.** Parameter estimates for the dose-response curves

| | $E_1$ | $\beta_1$ | $E_2$ | $\beta_2$ |
| --- | --- | --- | --- | --- |
| unrestricted | 1.01 | 0.49 | 1.50 | 0.99 |
| restricted | / | 0.49 | / | 2.79 |

**Fig 1.**
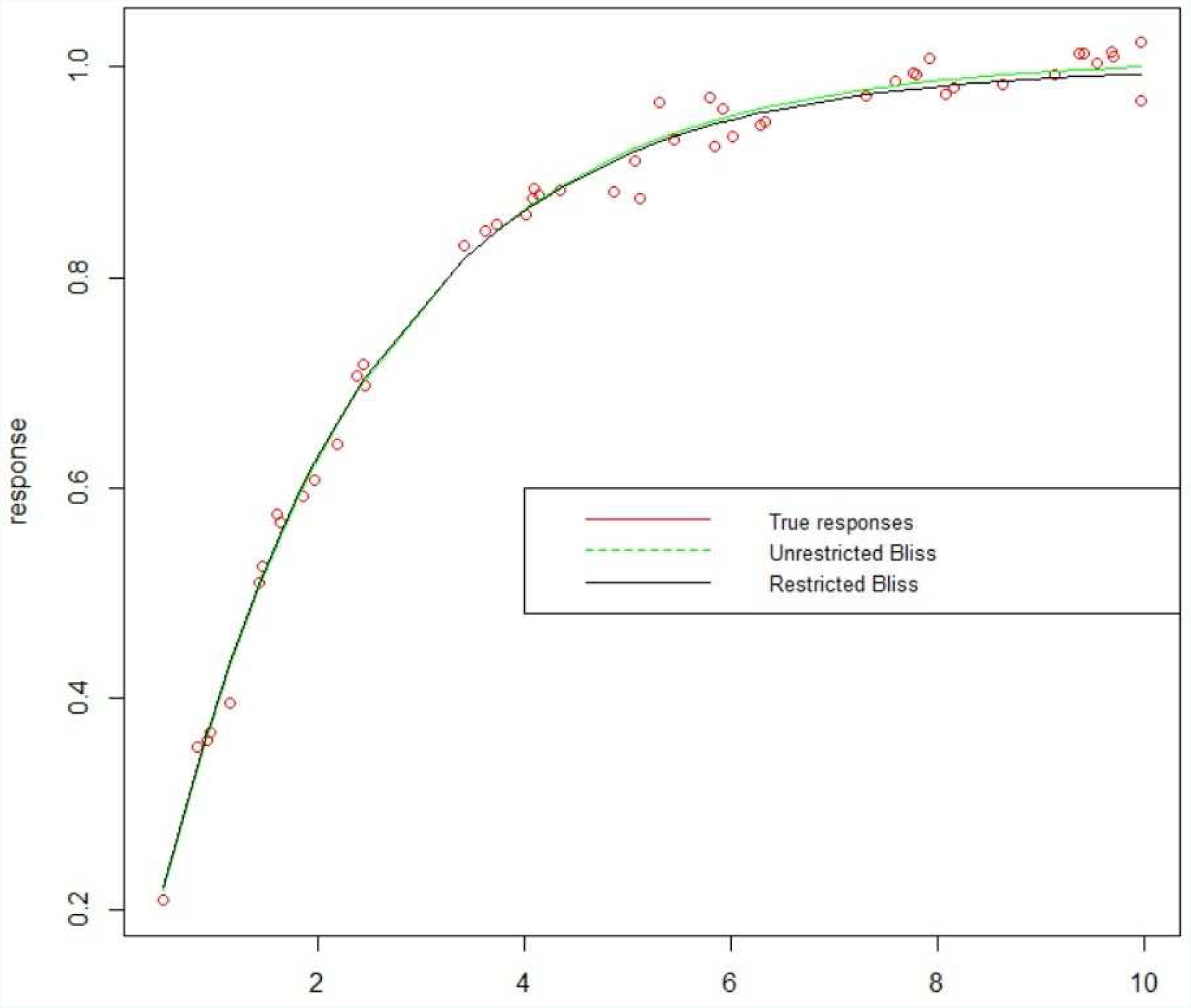
Estimated dose-response curves for the first data set

**Fig 2.**
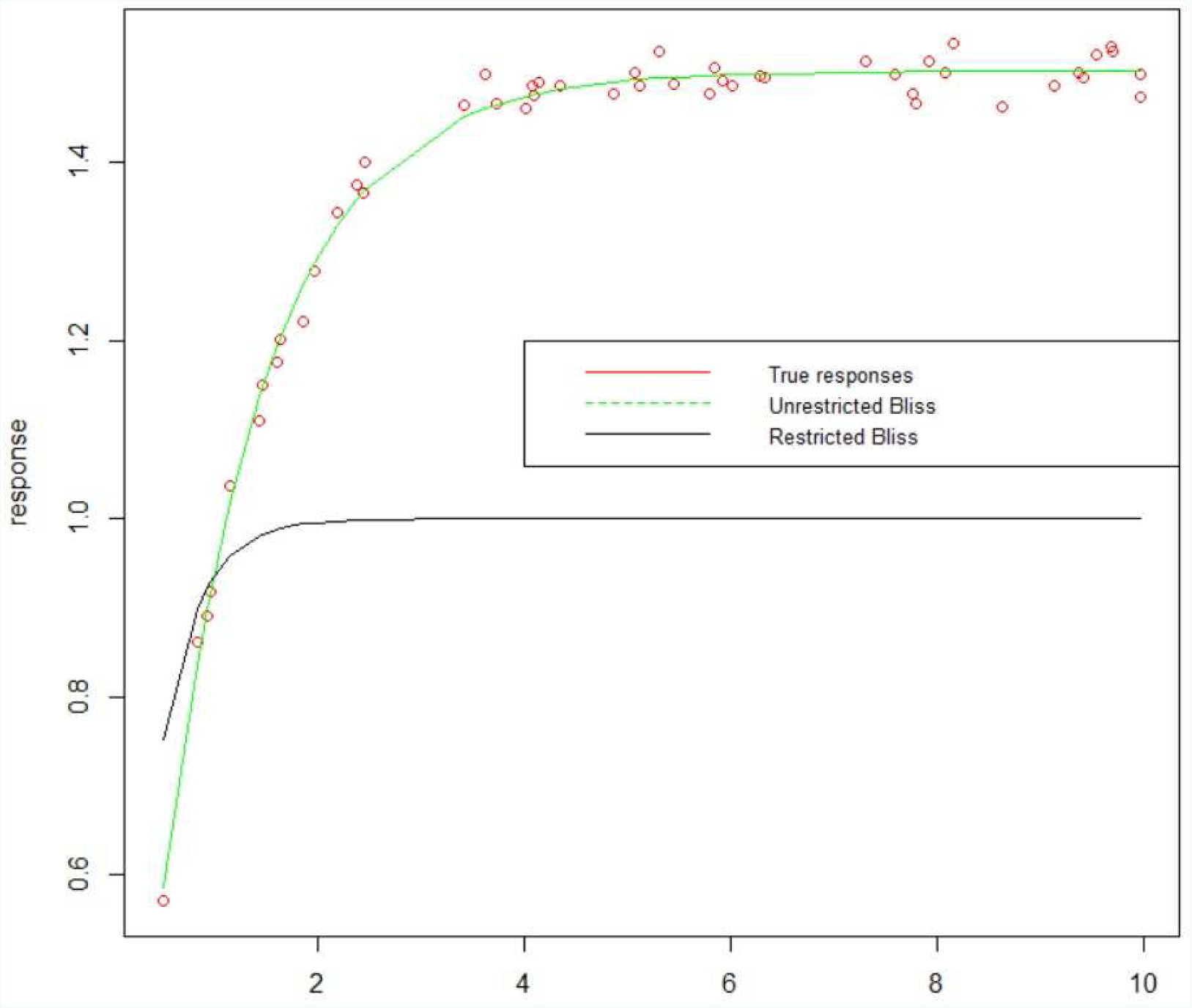
Estimated dose-response curves for the second data set

Finally, we compute the combined effects, as predicted by the restricted reference model and by the unrestricted reference model. These combined effects, denoted by 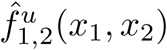 for the unrestricted reference model and by 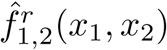 for the restricted reference model, are given by

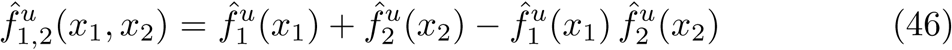

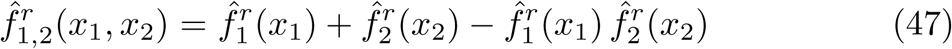

To illustrate the differences between both models, we compute the combined effects for every combination of doses *x*_1_ = 0.5, 1.0, 1.5, …, 10 and doses *x*_2_ = 0.5, 1.0, 1.5, …, 10, and this for both the restricted reference model and the unrestricted reference model. Next, we compute the relative differences *r*(*x*_1_, *x*_2_) between these combined effects, which we define as

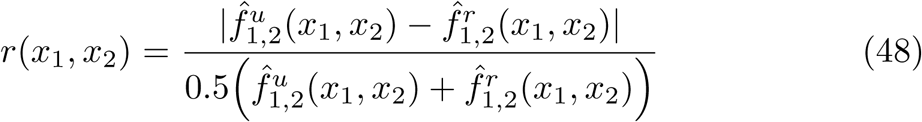

Fig. 3 shows *r*(*x*_1_, *x*_2_). It is seen that as long as both doses are small, the relative difference between 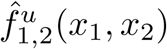 and 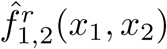 is also small, as the corresponding responses are far from the maximum effect. The relative difference is also small as *x*_1_ increases, because the first drug has maximum effect one, which is in correspondence with Bliss’ assumption. Furthermore, both 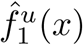 and 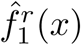 are very good approximations to the true first dose-response curve, as seen from Fig. 1, explaining the small difference between the estimated combined effects produced by both reference models. However, increasing *x*_2_ results in large differences in the predictions by both reference models. This is because the second drug has as maximum effect 1.5, although 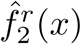 approximates the corresponding dose-response curve under the restriction that the maximum effect is 1. The larger the dose of the second drug, the closer the corresponding effect to the maximum effect, and the larger the difference between the restricted reference model and the unrestricted reference model. The consequences in a practical application might be severe, as the large differences in predicted combined effects imply that the interaction between two drugs might be evaluated incorrectly, for example by declaring that two drugs are synergistic while the opposite might be true.

**Fig 3.**
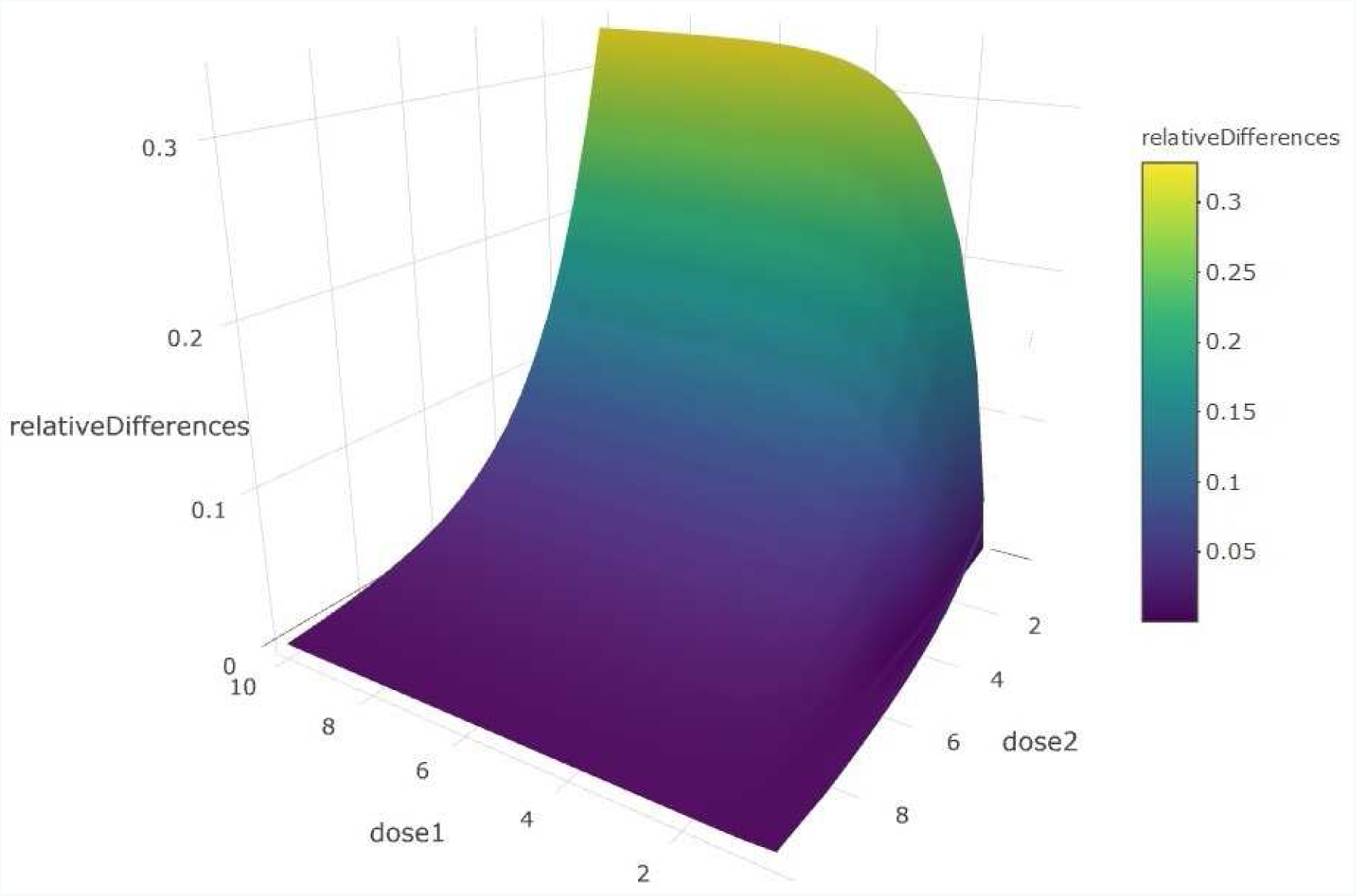
Relative differences between 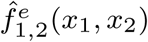 and 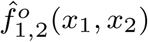

## 9 Conclusion

In this paper we have extended the Bliss independence model. The generalization is in two directions. First, the maximal effects of the considered single drugs are not assumed to be equal. Secondly, both the minimal effect, which is the same for all dose-response curves, as the maximal effects, are arbitrary. Furthermore, the constructed reference model applies to an arbitrary number of drugs. The model was obtained in several steps. First, two drugs are considered, and the corresponding dose-response curves are assumed to have the same maximal effect. By introducing a sound and elegant statistical similarity to the deterministic effect of a given dose of a given drug, we showed how the non-interaction principle introduced by Bliss can still be applied under less severe restrictions on the minimal and maximal effects. A general description of a reference model was given, relying on the idea, as adopted by the authors of the ZIP model, that a certain dose of the second drug is added to some dose of the first drug. We introduced two consistency principles that uniquely determine the unknown functions in this general description. In a second step the basic reference model was extended to apply to two drug that possibly have different maximal effects. Finally, the reference model was further generalized to apply to an arbitrary number of drugs. This general reference model is a genuine generalization of the reference model for two drugs, as it was constructed in terms of a recursion relation.

## Acknowledgements

This work was supported by the Norwegian Research Council project COLOSYS, under the frame of ERACoSysMed-1, the ERA-Net for Systems Medicine in clinical research and medical practice.

